# Bifocal tACS Enhances Visual Motion Discrimination by Modulating Phase Amplitude Coupling Between V1 and V5 Regions

**DOI:** 10.1101/2020.11.16.382267

**Authors:** Roberto F. Salamanca-Giron, Estelle Raffin, Sarah Bernardina Zandvliet, Martin Seeber, Christoph M. Michel, Paul Sauseng, Krystel R. Huxlin, Friedhelm C. Hummel

## Abstract

Visual motion discrimination involves reciprocal interactions in the alpha band between the primary visual cortex (V1) and the mediotemporal area (V5/MT). We investigated whether modulating alpha phase synchronization using individualized multisite transcranial alternating current stimulation (tACS) over V5 and V1 regions would improve motion discrimination. We tested 3 groups of healthy subjects: 1) an individualized In-Phase V1_alpha_-V5_alpha_ tACS (0° lag) group, 2) an individualized Anti-Phase V1_alpha_-V5_alpha_ tACS (180° lag) group and 3) a sham tACS group. Motion discrimination and EEG activity were compared before, during and after tACS. Performance significantly improved in the Anti-Phase group compared to that in the In-Phase group at 10 and 30 minutes after stimulation. This result could be explained by changes in bottom-up alpha-V1 gamma-V5 phase-amplitude coupling. Thus, Anti-Phase V1_alpha_-V5_alpha_ tACS might impose an optimal phase lag between stimulation sites due to the inherent speed of wave propagation, hereby supporting optimized neuronal communication.

**IMPACT STATEMENT:**

- Alpha multisite (V1 and V5) tACS influences global motion discrimination and integration
- Phase-amplitude coupling is associated with visual performance
- Multisite Anti-Phase stimulation of strategic visual areas (V1 and V5) is associated with connectivity changes in the visual cortex and thus, associated with changes in direction acuity

## INTRODUCTION

Motion direction discrimination training appears to be highly specific to the trained direction (Ball and Sekuler, 1987; Jia and Li, 2017) leading to the assumption that concurrent plastic changes may occur in early visual areas that are retinotopically organized and selective to basic visual features (Jehee et al., 2012; Karni and Sagi, 1991; Shibata et al., 2012). However, subsequent learning of a new direction is faster (Liu and Weinshall, 2000), suggesting the involvement of some higher-level processes. Furthermore, the manipulation of higher cognitive control processes, such as endogenous covert attention or exogenous spatial attention, improves stimuli location transfer and visual perceptual learning transfer to untrained stimulus location and features, respectively (Donovan et al., 2020; Donovan and Carrasco, 2018). Visual improvements would then rely on the interaction between multiple cortical areas (Gilbert et al., 2001), the combination of local intrinsic circuits and feedback connections from higher order cortical areas (Dosher and Lu, 1998; Gilbert and Sigman, 2007).

Specifically, research in humans (Blakemore and Campbell, 1969) and primates (Simoncelli and Heeger, 1998) has established that the primary visual cortex (V1) and medio-temporal areas (MT/V5, labeled henceforth as V5) are co-activated in complementary feedforward and feedback sweeps (Lamme and Roelfsema, 2000; Newsome and Pare, 1988), independent activation of these regions has been reported as well (Rodman et al., 1990). Their inter-dependency is related to the characteristics of the stimulus (e.g., orientation) and to the anatomical pathways that are recruited. Moreover, this channel is endowed with specific patterning of electrical signals in response to visuo-attentional perception, motion discrimination and memory encoding (Alagapan et al., 2019a; Polanía et al., 2012; Sauseng et al., 2009). In addition, evidence suggests that communication between these two regions in particular, may be established by phase synchronization of oscillations at lower frequencies (i.e., at Alpha-Beta frequencies, <25 Hz), acting as a temporal reference frame for information conveyed by high-frequency activity (at Gamma frequencies >40 Hz) (Bastos et al., 2015; Bonnefond et al., 2017; Fries, 2009; Seymour et al., 2019).

Phase synchronization is a key neuronal mechanism that drives spontaneous communication among dynamical nodes (Gollo et al., 2014), implying that this mechanism supports attentional, executive, and contextual functions (Doesburg et al., 2009; Freunberger et al., 2007; Palva and Palva, 2011). The two simplest phase synchronization patterns are *in-phase synchronization* (i.e., zero phase lag between the two regions) and *anti-phase synchronization* (i.e., 180° phase lag between the two regions). In-Phase synchronization between two distant neuronal populations is thought to subserve the integration of separated functions that are performed in these different regions (Engel et al., 1991; Roelfsema et al., 1997; Wang et al., 2010). Conversely, anti-phase patterns reflect more dynamical reciprocity, where certain areas of the brain increase their activity while others decrease their own activity. Such anti-phase patterns have been reported during sleep (Horovitz et al., 2009), or during visual attentional tasks (Yaple and Vakhrushev, 2018). It has been proposed that these anti-phase oscillation patterns reflect time-delays in functional coupling between two connected regions (Petkoski and Jirsa, 2019). Since communication between neurons is achieved by propagation of action potentials throughout axons, with conduction times defined by some regional specificities, such as myelination density, number of synaptic relays, inhibitory couplings etc., an optimal phase delay relationship between two interconnected regions could be a key driver of brain communication.

In this article, we set out to determine whether motion discrimination performance can be enhanced when ‘artificially’ entraining/manipulating the phase relationship between V1 and V5. This is based on the idea that inter-areal synchronization plays a significant role in V1-V5 communication, as demonstrated previously (Lewis et al., 2016; Siegel et al., 2008). We used individually adjusted, Alpha transcranial alternating current stimulation (tACS) to entrain endogenous oscillations (Helfrich et al., 2014) and enhance inter-areal information flow (Zhang et al., 2019). The modulation consisted in applying approximately 15 minutes of concurrent, bifocal (over V1 and V5), individualized Alpha-tACS. We assessed two conditions of stimulation: In-Phase (zero phase lag) stimulation and Anti-Phase stimulation (180° phase lag). This was done to contrast the behavioral consequences of these two different phase delays (Klimesch et al., 2007). A Sham tACS group was evaluated to control for non-specific, placebo-like effects.

Furthermore, the entire experiment was conducted while recording multi-channel electroencephalography (EEG). Electrophysiological analyses were computed with the objective of determining EEG markers of interareal modulation between the two target areas. We paid special attention to connectivity metrics in the Alpha band, as well as in the Gamma band because of their role in visual features binding (Elliott and Müller, 1998; Gray and Singer, 1989; Zhang et al., 2019). In fact, the interactions between Alpha and Gamma oscillations may serve as a framework supporting the feedforward and feedback loops of inter-regional brain communication within the visual system (Kerkoerle et al., 2014; Michalareas et al., 2016). Specifically, top-down Alpha appears to control the timing and elicitation of higher frequency rhythms, thus optimizing communication in the visual cortex (Fries, 2015; Michalareas et al., 2016). Taken together, we hypothesize that the best inter-areal Alpha phase relationship for optimal oscillatory entrainment leading to respective behavioral enhancement is associated with changes in Alpha-Gamma coupling.

## RESULTS

All participants tolerated the stimulation well and did not report any adverse effects such as peripheral sensory or phosphene perception. Five participants could not be included in the analyses: One participant discontinued the experiment without stating the reason for it and four participants were discarded, because they failed to perform the task properly. Therefore, 45 full sets of data were analyzed, forming homogenous groups of 15 participants. For the EEG metrics of interest (ZPAC), three data points (i.e., 2 from the In-Phase group, 1 from the Anti-Phase group) were found by Cook’s Distance algorithm (Cook, 1977) to be more than two standard deviations from the mean of the distribution, and were thus not included in the analyses.

### Motion direction performance throughout groups and time

Figure 2A displays the mean baseline-corrected NDR thresholds across participants, reflecting the normalized motion direction value corresponding to 75% correct performance (see Method section) across groups and time. Although there was no statistically significant difference between groups at baseline (Anti Phase vs. In Phase b = 1.670, P = 0.809, CI = −11.835 15.175, Sham vs. In Phase b = 3.260, P = 0.624, CI = −9.770 16.290, Sham vs Anti Phase b = 1.590, P = 0.815, CI = −11.696 14.876; see also Supplementary Table 1 providing the raw NDR values), the baseline values showed a large variability, therefore we applied a baseline correction procedure to account for this variability. When considering all the groups together, the change in baseline-corrected NDR was not significant between TP0 and TP10 (b = −0.05, P = 0.189, CI = −0.124 0.024) nor between TP0 and TP30 (b = −0.067, P = 0.079, CI = −0.141 0.008), neither between TP10 and TP30 (b = −0.017, P = 0.657, CI = −0.091 0.057). However, there was a significant difference at TP0, TP10 and TP30 between the In-Phase and the Anti-Phase group (b = 0.257, P = 0.015, CI = 0.05 0.464). There was no difference for other group comparisons for all time points (b = 0.16, P = 0.118, CI = −0.04 0.36 Sham and In-Phase; b = −0.097, P = 0.349, CI = −0.301 0.107 Sham and Anti-Phase).

For the Anti-Phase group the changes in the NDR were not significant between Baseline and TP0 (b=−2.865, P=0.31, CI=−8.401 2.671), however they were strongly significant between Baseline and TP10 (b=−9.655, P=0.001, CI=−15.19 −4.119) and between Baseline and TP30 (b=−14.519, P=0.001, CI=−20.054 −8.983). Moreover, NDR was significant between TP0 and TP10 (b=−6.79, P=0.016, CI=−12.325 −1.254), and between TP0 and TP30 (b=−11.653, P>0, CI=−17.189 −6.118), although not significant between TP10 and TP30 (b=−4.864, P=0.085, CI=−10.4, 0.672).

For the In-Phase group the changes in the NDR were not significant between Baseline and TP0 (b=0.23, P=0.93, CI=−4.881 5.342), nor between Baseline and TP10 (b=− 2.309 P=0.376, CI=−7.42 2.802) neither between Baseline and TP30 (b=−0.291, P=0.911, CI=−5.403 4.82). Moreover, NDR was not significant between TP0 and TP10 (b=−2.539, P=0.33, CI=−7.65, 2.572), nor between TP0 and TP30 (b=−0.522, P=0.841, CI=−5.633 4.59), neither between TP10 and TP30 (b=2.017, P=0.439, CI=−3.094, 7.129).

For the Sham group the changes in the NDR were marginaly significant between Baseline and TP0 (b=−5.802, P=0.04, CI=−11.339 −0.265) and between Baseline and TP30 (b=−6.311, P=0.025, CI=−11.849 −0.774), but not significant between Baseline and TP10 (b=− 4.577 P=0.105, CI=−10.114 0.96). Moreover, NDR was not significant between TP0 and TP10 (b=1.225, P=0.665, CI=−4.312 6.762), nor between TP0 and TP30 (b=−0.509, P=0.857, CI=−6.047 5.028), neither between TP10 and TP30 (b=−1.734, P=0.539, CI=−7.272 3.803).

**Figure 2.**
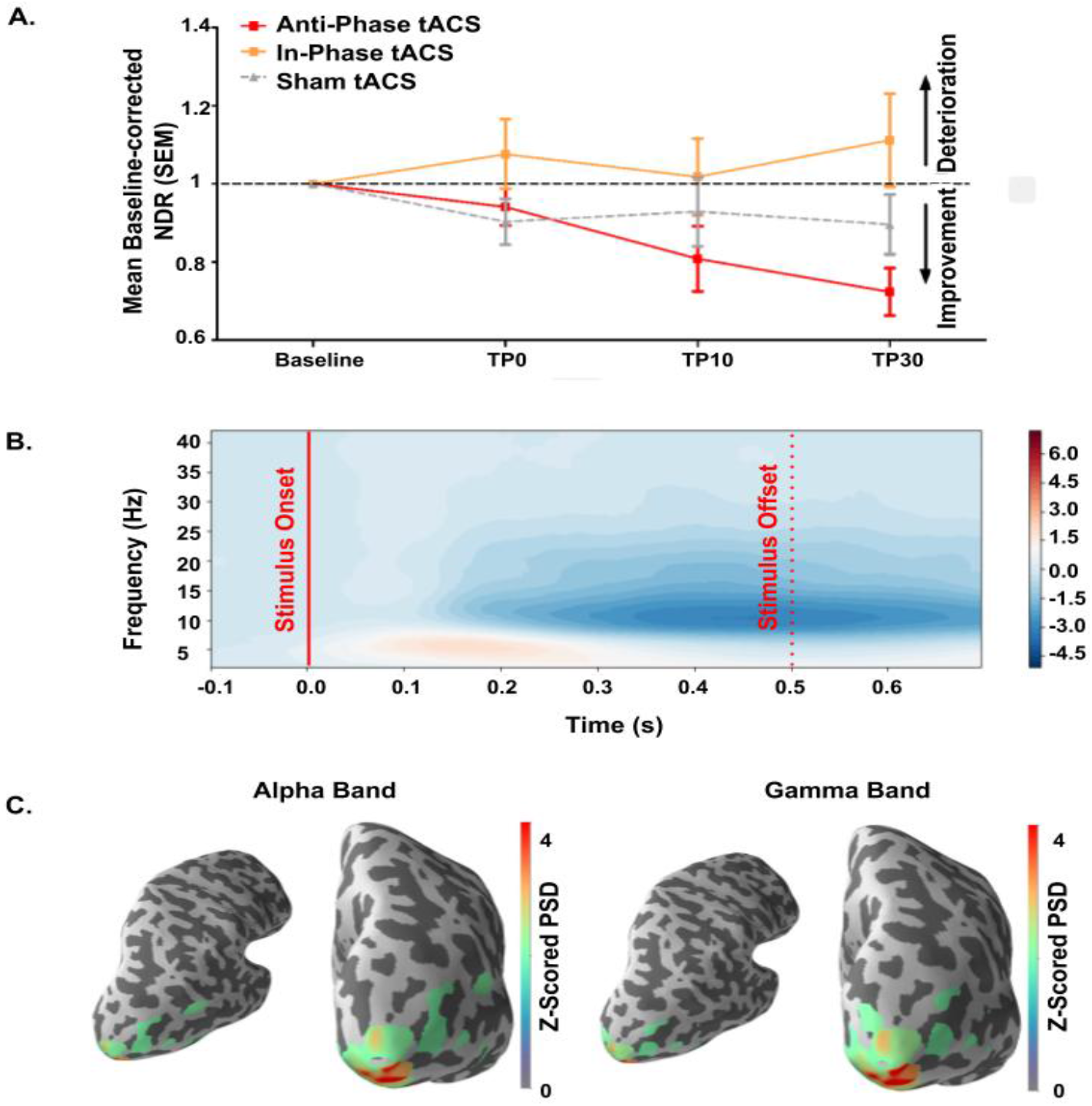
**(A) Baseline-corrected NDR (Normalized Direction Range) threshold** evolution across time-points for the three stimulation conditions. Bars correspond to Standard Errors of the Mean (SEM). Anti-Phase stimulation induced an increased performance translating into a significantly pronounced behavioral improvement over time at the group level. The behavioral performance of the Anti-Phase group was significantly enhanced compared to the In-Phase group. **(B) Time-frequency representation** of the averaged response during a trial at the baseline period, before the stimulation. It shows a typical Event Related Synchronization at (ERS) the Theta/Alpha band, followed by an Event Related Desynchronization (ERD) in the Beta band. (C) **PSD projected on 3D brain**.

### EEG Results

In all participants, the visual discrimination task led to an amplitude increase in the Theta/Low Alpha band, right after the onset of the stimulus, followed by a phasic decrease in power in the High Alpha/Low Beta bands ~200 ms thereafter (**Figure 2B**). Additionally, in frequencies above 30 Hz, there was a constant decrease in magnitude during stimulus presentation, as previously described in the literature for this type of visual task (e.g., (Siegel et al., 2007; Townsend et al., 2017)).

The Lasso model, defined for each time point, showed that a single EEG marker, namely ZPAC-V1p_Alpha_V5a_Gamma_ had the largest explanatory value for the variance of NDR at TP10 (R^2^=0.057, λ=0.114) and TP30 (R^2^=0.082 λ=0.052), irrespective of the stimulation group.

Since the ZPAC-V1p_Alpha_V5a_Gamma_ values best explained changes in the performance after stimulation, the rest of the manuscript focuses on this metric in order to further explore stimulation and time effects. The opposite direction, ZPAC-V1a_Gamma_V5p_Alpha_ was used as a control analysis to test for the directional specificity of the present results.

### Changes in bottom-up V1 Alpha phase (V1p_Alpha_) - V5 Gamma amplitude (V5a_Gamma_) coupling

**Figure 3A** shows the mean baseline-corrected ZPAC-V1p_Alpha_V5a_Gamma_ values for the three groups across time. These values were extracted from the significant modulation of interest between the Alpha/High Theta and the Low Gamma bands shown in Figure 3B. It reveals a significant diminishment in the Alpha/High Theta (5-12 Hz) – Low Gamma (30-42 Hz) phase amplitude coupling at TP10 for the Anti-Phase and the Sham group and a significant augmentation in coupling for the In-Phase group. At TP30, there is overall a more prominent augmentation of the coupling for the In-Phase group, a more pronounced diminishment for the Anti-Phase and rather a stable response for the Sham group. To statistically analyze the descriptive differences between the three conditions, we computed a mixed linear model on the ZPAC-V1p_Alpha_V5a_Gamma_values. The model returned a marginally significant change over time between the interval TP10 and TP30 (b = −0.769, P = 0.055, CI = −1.556, 0.018), but no significant differences between the Anti-Phase and the In-Phase groups (b = 0.836, P = 0.35, CI =−0.916, 2.588). This held true also when comparing the Anti-Phase and Sham groups (b = 1.009, P = 0.249, CI = −0.708 2.726), and the In-Phase and Sham groups (b = 0.173, P = 0.84, CI = −1.51 1.856).

When ZPAC-V1p_Alpha_V5a_Gamma_values were entered as a single confounder into the baseline-corrected NDR model, it did not significantly account for the overall variance for all the stimulation groups at all time points (b = 0.015, P = 0.196, CI = −0.008, 0.039). However, ZPAC-V1pV5a from the Anti-Phase group as compared to the In-Phase group, did significantly account for the variability of the NDR as a fixed effect over time at both TP10 and TP30 (b = 0.071, P = 0.048, CI = 0.001, 0.142). This was not the case when comparing the ZPAC-V1pV5a values from the In-Phase group *versus* those from Sham (b = −0.023, P = 0.44, CI = 0.081, 0.035), nor when comparing those from Anti-Phase and Sham groups (b = 0.048, P = 0.095, CI = −0.008, 0.105) at any of the two time points (all other comparisons are shown in the Supplementary Table 2).

**Figure 3.**
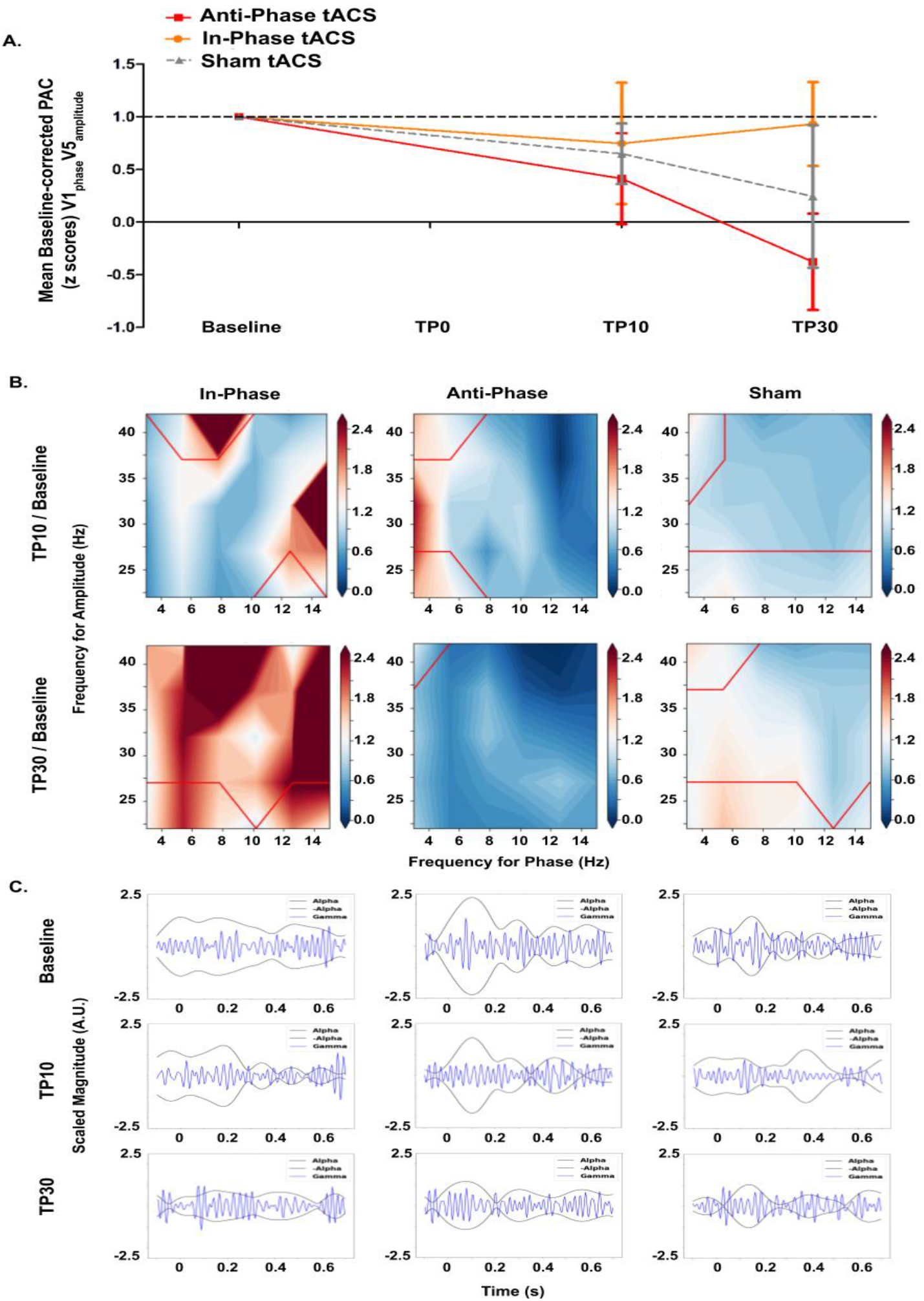
**(A) Baseline-corrected, bottom-up V1-Alpha phase V5-Gamma Amplitude coupling across time-points**. Bars correspond to Standard Errors of the Mean (SEM). Please note the strong decrease for the In-Phase group towards TP30. (**B) Averaged, baseline-corrected, significant clusters (p<0.5) from the V1-Gamma amplitude V5-Alpha phase coupling spectrums** for the three stimulation groups and for the two time points after stimulation averaged during the stimulus presentation interval. **(C) Alpha V1 Gamma V5 Phase-amplitude coupling during stimulus presentation**

### Changes in top-down V1 Gamma amplitude (V1a_Gamma_) - V5 Alpha phase (V5p_Alpha_) coupling

To test the eventual directional specificity of the present results, we examined the opposite phase-amplitude coupling between V1 and V5. **Figure 4A** provides the descriptive data for the ZPAC-V1a_Gamma_V5p_Alpha_ for all 3 experimental groups over time. To statistically analyze these data, we applied a comparable approach as in the previous section. **Figure 4B** shows the results for the ZPAC-V1a_Gamma_V5p_Alpha_, which appeared to have a significant Alpha/Theta – Low Gamma phase amplitude cluster at both TP10 and TP30. Diminished coupling is evident for the three stimulation groups when V5 Alpha/ High Theta (6-10 Hz) modulated Low V1 Low Gamma (30–37 Hz) amplitude. We then built a similar mixed linear model using the ZPAC-V1a_Gamma_V5p_Alpha_ values. These analyses showed no significant change in time between TP10 and TP30 (b = 0.409, P = 0.286, CI = −0.343, 1.161). Neither at TP10 nor at TP30 was a significant difference between the Anti-Phase and Sham group (b = −0.718, P = 0.484, CI = −2.727, 1.292), between the Anti-Phase and In-Phase group (b = 0.695, P = 0.506, CI = −1.353, 2.744) or between the In-Phase and Sham group (b = − 1.413, P = 0.161, CI = −3.39, 0.564). Unsurprisingly, when ZPAC-V1a_Gamma_V5p_Alpha_ was entered as a confounder into the NDR model, it did not significantly account for the variance in NDR scores for all the stimulation groups together at all time points (b = −0.007, P = 0.53, CI = −0.029, 0.015). Additionally, there was an absence of a significant interaction between ZPAC-V1a_Gamma_V5p_Alpha_ and each stimulation group, suggesting that the ZPAC-V1a_Gamma_V5p_Alpha_ group values did not explain the group differences in the NDR values at all timepoints (In-Phase vs. Anti-Phase: b = −0.055, P = 0.432, CI = −0.191, 0.082, In-Phase vs. Sham: b = 0.006, P = 0.908, CI = −0.09, 0.101, Anti-Phase vs. Sham: b = 0.06, P = 0.234, CI = −0.039, 0.16) (all other comparisons are shown in the Supplementary Table 3).

**Figure 4.**
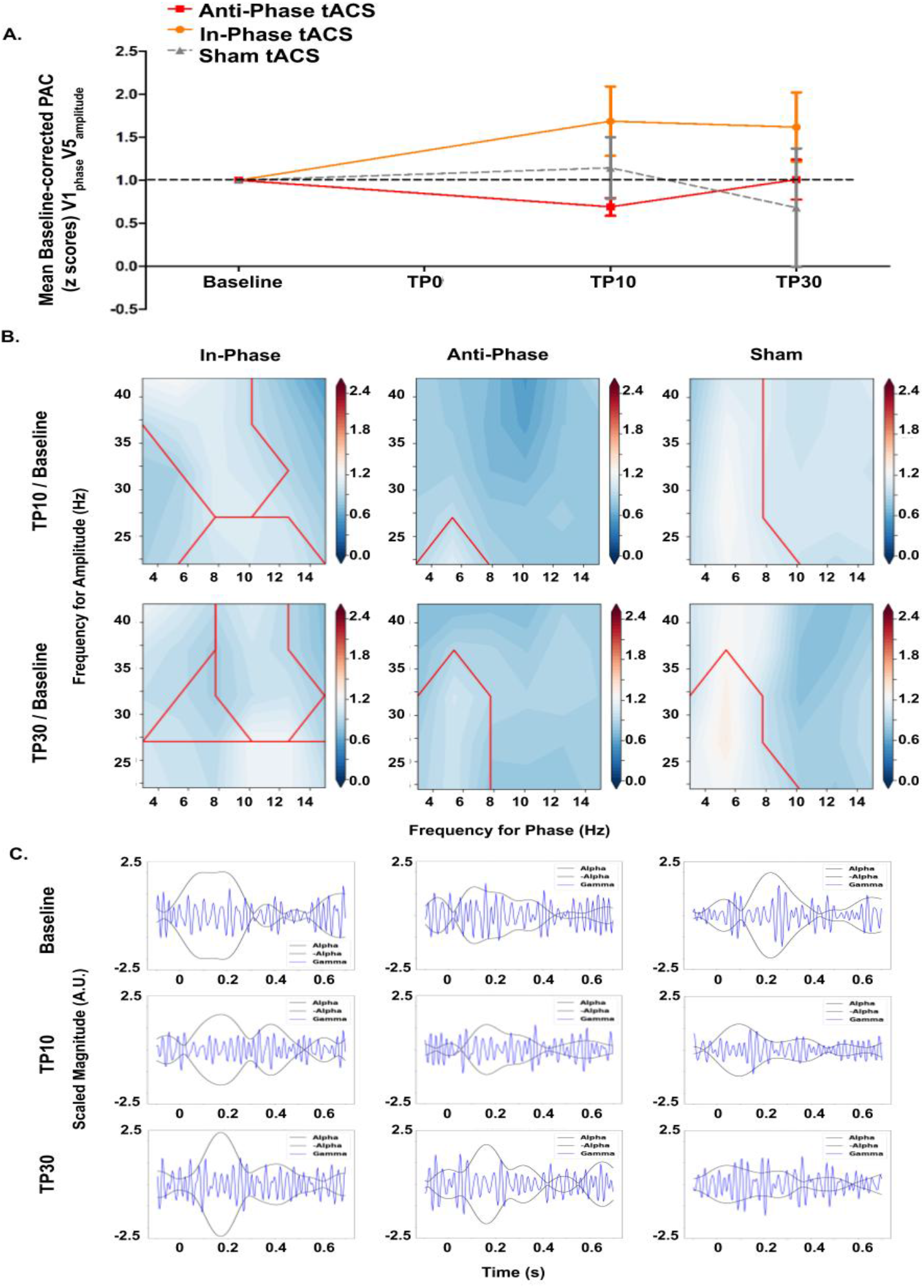
**(A) Baseline-corrected, top-down V1-Gamma amplitude V5-Alpha phase coupling across time-points.** Bars correspond to Standard Errors of the Mean (SEM). Please note the strong decrease for the In-Phase group towards TP30. **(B) Averaged, baseline-corrected, significant clusters (p<0.5) from the V1-Gamma amplitude V5-Alpha phase coupling spectrums** for the three stimulation groups and for the two time points after stimulation averaged during the stimulus presentation interval. **(C) Gamma V1 Alpha V5 Phase-amplitude coupling during stimulus presentation**

## DISCUSSION

By applying multisite tACS in the Alpha range to V1 and V5 with a phase difference of 180 degrees (Anti-Phase) during a visual global motion direction discrimination and integration task, we were able to modulate interactions between V1 and V5 functionally-relevant resulting in significant behavioral improvement. For instance, this led to a significant enhancement of motion discrimination and integration in the young healthy individuals. Specifically, the behavioral improvement was associated with a modulation of inter-regional oscillatory coupling between the two stimulated brain areas. The three main findings can be summarized as follows: 1) *Anti-Phase* V1_Alpha_-V5_Alpha_ tACS entrainment leads to an improvement in visual performance during and shortly after stimulation compared to *In-Phase* V1_Alpha_-V5_Alpha_, which appears rather detrimental to motion discrimination and integration, 2) improved performance with *Anti-Phase* V1_Alpha_-V5_Alpha_ tACS can best be explained by changes in bottom-up V1 Alpha phase - V5 Gamma amplitude coupling (*ZPAC-V1p_Alpha_V5a_Gamma_*), and 3) the opposite, top-down modulation (*ZPAC-V5p_Alpha_V1a_Gamma_*) did not influence performance in the current paradigm.

### In-Phase V1_Alpha_-V5_Alpha_ stimulation hampers motion discrimination and integration

In-Phase tACS between two distant regions is motivated by the idea of increasing interregional synchronization and connectivity within a network (Polanía et al., 2012; Schwab et al., 2019; Vieira et al., 2020), under the hypothesis that a reduced phase-lag (~0°) between sites would promote an optimal inter-areal coupling and thus, optimal communication (e.g., (Fries, 2005)). There is empirical evidence supporting this hypothesis. For instance, In-Phase stimulation has been associated with increased performance in visuo-attentional and memory tasks (Alagapan et al., 2019b; Polanía et al., 2012; Violante et al., 2017), together with increased phase synchronization in the stimulated frequency band. In contrast to these data however, the present results showed opposite effects, i.e. the In-Phase condition rather impaired visual discrimination capacity during the stimulation period of ~13±2 minutes, and performance did not improve, but rather decreased 10 and even 30 minutes after applying it.

Visual discrimination is associated with local Alpha desynchronization right after stimulus presentation (Dijk et al., 2008; Erickson et al., 2019; Hillyard et al., 1998; Sauseng et al., 2009; Zammit et al., 2018). Subsequently, it has been shown in several perceptual experimental modalities that a decrease in the Alpha-Beta band is linked to better stimulus perception (Griffiths et al., 2019). Thus, a high amplitude and zero-phase lag condition might not be optimal in this case because, as shown in the present data, an increased V1 Alpha phase - V5 Gamma amplitude coupling post stimulation is rather associated with poor performance. It might be an intricated orchestration of oscillatory signatures that travels throughout the clusters of the neural network, controlled by stimuli properties (Muller et al., 2018). This oscillatory orchestration could be modeled as a multi-level interacting dynamical system (Alexander et al., 2019). Ultimately, cognition relies on feedback and feedforward dynamics, and these processes are only possible through complex, well-orchestrated phase and amplitude interactions (Siegel et al., 2012).

From a more integrative perspective, the inhibition timing hypothesis (Klimesch, 2012) states that the optimal electrophysiological scenario that promotes perception relies on an inter-regional interplay of Alpha inhibition and Alpha disinhibition among areas belonging to the same network, as shown in the visual cortex (Shen et al., 2011). When this precise timing of activation/deactivation is disrupted by enforced Alpha In-Phase rhythms, it might generate a subsequent flood of massively synchronized signals, creating an artificial source of noise that may prevent accurate perception of stimulus features (Faisal et al., 2008; Voytek and Knight, 2015). Hence, the neuronal oscillatory system might require some time to come back to its basal processing state, just as demonstrated for the overall performance at 10 min and 30 min after stimulation. Additionally, although the noise created by the In-Phase synchronization could be beneficial under some stochastic resonance phenomena (Wiesenfeld and Moss, 1995), because of its randomness nature, it harms the idea of an ordered, well-defined oscillatory gating process. This gating process ought to include specific frequency signatures between network pathways and clear time-space streams of activity (Jensen et al., 2014; Richter et al., 2017), instead of an equal probability of appearance of several frequency components across time without following a master order, characteristic of stochastic circumstances.

Furthermore, one could expect an energy optimization over time of the Alpha oscillations in the visual cortex under the Hamiltonian premise of minimum action in electrically charged natural systems (Aitchison and Lengyel, 2016; Seung et al., 1998). The lower the energy, the less prominent the power trace is, resulting in turn, into a weaker synchronization between the two signals. In other words, a fewer demand of resources and a less complex gating operation tends to a more prominent oscillatory desynchronization trace (Bays et al., 2015).

In conclusion, positive behavioral effects are not necessarily associated with an In-Phase synchronized magnification of the Alpha occipital rhythms in a visual discrimination task, but rather an ordered gating of oscillations and patterns as the Anti-Phase condition promotes.

### Anti-Phase V1_Alpha_-V5_Alpha_ stimulation enhances motion discrimination and integration

The improved offline performance reported in the present study is in accordance with a body of literature showing that inter-areal Anti-Phase stimulation might boost behavior in several contexts. For instance, Beta band Anti-Phase bi-hemispheric stimulation has been shown to increase visual attentional capacity (Yaple and Vakhrushev, 2018). In the same vein, Theta band Anti-Phase stimulation over the prefrontal and perysilvian area has been found to improve controlled memory retrieval (Marko et al., 2019), while Gamma band Anti-Phase stimulation between the cerebellum and M1 enhances visuomotor control (Miyaguchi et al., 2019). Here, we found that Anti-Phase V1_Alpha_-V5_Alpha_ tACS applied on average for 13±2 minutes during a motion discrimination task significantly boosted motion direction discrimination and integration 10 minutes after the end of the stimulation and the effects continued to strengthen even 30 minutes later.

While any after-effects of tACS are under debate in the field (Strüber et al., 2015), we think that the improved performance measured in the Anti-Phase group, which persists over time, are not simply explained by an offline effect of the stimulation per se. Instead, we argue that it is the repeated practice of the task combined with the Anti-Phase tACS condition that promotes a “learning-like after-effect”. These after-effects might indeed find a justification in the accumulation of offline effects that lead to a carry-over of the achieved behavioral improvement (Heise et al., 2019) and might generate favorable plastic changes in the visual cortex due to the learning associated with the task, as it has been shown in non-human primates (Yang and Maunsell, 2004).

The biophysical mechanisms underlying the behavioural improvement, as well as its relative timing are still unclear. One can speculate that Alpha oscillatory traces should be considered as traveling flows of electrical activity around the specific neuronal network (Alamia and VanRullen, 2019; Lozano-Soldevilla and VanRullen, 2019), instead of simple mono-focal fluctuating rhythms. Under this premise, at really specific timings, these waves are used to either start or stop inhibition inter-regionally with the objective of pursuing an optimal transmission of stimulus information, and more importantly, a sustained perceptual learning (Sigala et al., 2014).

Alekseichuk and colleagues compared intracranial recordings in the temporal area of macaques undergoing frontoparietal 10Hz Anti-Phase or In-Phase stimulation as well as the voltage and electric field distribution associated with the two stimulation modes (Alekseichuk et al., 2019). Results showed a higher electric field magnitude, plus an unidirectional concentration of field lines for the Anti-Phase condition, whereas for the In-Phase condition there was a reduced magnitude and a bidirectional flow of electric field lines. The present electrical field simulation globally revealed similar spatial patterns suggesting that Anti-Phase stimulation generates more dynamical changes in electrical field distribution, resembling the traveling wave phenomenon with specific dynamics across time and characterized by a specific propagation speed. Alpha-band travelling waves recorded with EEG under stimulus-driven conditions are being increasingly investigated (e.g., (Hindriks et al., 2014; Lozano-Soldevilla and VanRullen, 2019). A more accurate description of travelling waves, especially between V1 and V5 areas, which are relatively close, would require a multi-modal imaging approach combining high temporal and spatial resolution (Giannini et al., 2018). However, using EEG-derived phase amplitude coupling, it is possible to infer directionality of signal flow (Nandi et al., 2019). The direction of the coupling is assumed to be bottom-up if the modulating signal (Alpha band) is recorded in a primary functional neuronal population, located in lower anatomical areas (V1) whereas the carrier signal (Gamma band) is rather on higher cognitive and anatomical areas (MT/V5), receiving inputs mainly from other regions of the cortex (Jiang et al., 2015). Otherwise, the interaction ought to be top-down. This finds justification from a signal processing perspective as well, where the power of the carrier signal is being modified under the phase of the modulating signal.

Visual stimulus onset has been shown to trigger propagating rhythms in the primary and secondary visual cortices of monkeys, leading to a specific phase relationship between the oscillations at both sites (Muller et al., 2014). In humans, propagation of feedforward flows have been reported during visual motion discrimination, with latencies modulated by characteristics of the stimulus (Sato et al., 2012; Seriès et al., 2002). Moreover, traveling waves in the posterior cortex measured by intracortical recordings, show a modulation of Gamma amplitude through Alpha phase control, with velocities among 0.7-2 m/s (Bahramisharif et al., 2013), corresponding approximately to half a cycle of an Alpha band oscillation. Then, this half Alpha phase-lag between stimulation sites, induced by the Anti-Phase condition, could aid neuronal communication, because of the inherent speed of propagation of the signals.

### Changes in bottom-up V1-Alpha phase - V5-Gamma amplitude coupling, but not the opposite direction, explain improved performances induced by Anti-Phase V1_Alpha_-V5_Alpha_ stimulation

The present positive behavioral effects were associated with a bottom-up V1-Alpha phase V5-Gamma amplitude decrease in coupling. This measure reflects the idea that the feedforward direction between V1 and V5 is regulated by a controlled amplitude modulation of Alpha-V1 over the phase of Gamma-V5, which scales with improved motion discrimination in the Anti-Phase group. This suggests the idea that there is an optimal range of Alpha rhythm magnitude that is more favorable to generate trains of local Gamma bursts, which might convey the most relevant information of the visual stimulus’ features to promote motion discrimination (Nelli et al., 2017; Tu et al., 2016).

This bottom-up Alpha-Gamma interaction is in line with the theory of cross-frequency nested oscillations (Bonnefond et al., 2017). Accordingly, the organization of tasks in the visual system is done through the timed gating of information encoded in local Gamma bursts, happening every 10-30 ms and that are regulated through the Alpha inhibitory role (Jensen et al., 2014). Additionally, our finding that changes in phase amplitude coupling between Alpha-V1 and Gamma-V5 predict behavioural improvements in the Anti-Phase group is congruent with the fact that motion discrimination has been shown to occur as a feedforward oscillatory phenomenon (Seriès et al., 2002), and that these oscillations in the occipital cortex do not only belong to a single frequency band, but rather to a modulation of Alpha and Gamma rhythms (Bahramisharif et al., 2013).

Thus, not only Alpha (Alamia and VanRullen, 2019), but also Gamma oscillations in the visual cortex appear as phase-sensitive propagating waves following maximal flow of information (Besserve et al., 2015). Alpha activity as the idling interareal rhythm of the brain, typically gets perturbed, when there are local bottom-up inputs (von Stein et al., 2000). Bottom-up inputs that become evident as Gamma activity carrying novelty of a stimulus (Gray, 1999, p. 199). This mechanism has been reported to be linked to plastic changes in the visual system (Gray, 1999, p. 199), in the same way as we had hypothesized: it occurs in the present Anti-Phase stimulation, likewise associated to the bottom-up flow of information processing.

Finally, we did not find any significant changes in the opposite top-down V5-Alpha phase - V1-Gamma Amplitude coupling and the values measured 10 minutes and 30 minutes after stimulation did not account for changes in motion discrimination performance or their variance. Although recordings in monkeys’ visual cortex have shown a top down Alpha-Beta that granger-causes a bottom-up Gamma rhythm (Richter et al., 2017), it does not necessarily contradict our findings since what we report reflect bottom-up coupled nested oscillations from one neuronal cluster to another, rather than a causal generation of oscillatory activity from one site to another. These markers indeed imply two different processes of interaction, in most of the circumstances mutually exclusive. Then, there might be different cross-frequency mechanisms that sustain visual discrimination that are revealed by these different electrophysiological markers. Exploring this variety of markers might lead to a better understanding of neural communication supporting visual discrimination.

## CONCLUSIONS

The present experiments revealed that entraining the organization of Anti-Phase oscillation patterns between V1 and V5 during motion discrimination using bi-focal tACS can enhance performance persisting even after the stimulation period. These after-effects were mechanistically partially explained by changes in bottom-up V1-Alpha V5-Gamma Phase-Amplitude coupling, while the inverse direction did not play any significant role at explaining the behavioral performance. These new results might be explained by the concept of traveling waves from V1 to higher visual areas, as well as the precise phase-timing hypothesis. It is indeed likely that an optimal phase-lag between stimulation sites, induced by the Anti-Phase tACS entrainment, did promote neuronal communication because of the inherent speed of wave propagation. Furthermore, we could infer that Alpha Anti-Phase stimulation, acts as a controller of the Alpha disinhibition-gating capacities and as such, modulates bottom-up trains of Gamma bursts in the V1-V5 pathway. The precise characteristics of the Gamma bursts (e.g., phase, time) might play a significant role in improving the performance in motion discrimination.

The present findings point towards the the exciting potential of the current approach to be extended towards an ameliorated stimulation orchestration with cross-frequency montages targeting the motion discrimination pathway. Furthermore, it potentially opens a novel direction of non-invasive interventions to treat patients with deficits in the visual domain, such as after a stroke.

## MATERIALS AND METHODS

### Subjects

50 healthy subjects were recruited (range age: 18 to 40 years old, 24 females). All individuals were right handed with normal or corrected to normal vision, and had no history of neurological diseases or cognitive disability. A written consent form was obtained from all participants prior the experiment. The study was performed according to the guidelines of the Declaration of Helsinki and approved by the local Swiss Ethics Committee (2017-01761).

### Study design

Individual testing started with a familiarization phase followed by the actual experiment. During the familiarization phase, we ensured that the subject understood the visual discrimination task and reached stable performance. After EEG acquisition was prepared, a baseline block, which consisted of a task-related EEG recording without tACS was started. After a few minutes of rest, electrodes were placed over the occipital and temporal cortex, and electrical stimulation was started, remaining on for the entire duration of the block. Immediately after the start of stimulation, the second timepoint (TP0) was recorded with concurrently-measured EEG. Thereafter, the stimulation electrodes were removed and after a few minutes of rest, two succeeding evaluation points (TP10: 10 minutes after stimulation, TP30: 30 minutes after stimulation) were measured using the same task-related EEG setup, without tACS (see **Figure 1A**).

### Visual discrimination task

The visual task used is a well-established 2-alternatives, forced-choice, left-right, global direction discrimination and integration task (150 trials per time point) (Das et al., 2014; Huxlin et al., 2009). The stimulus consisted of a group of black dots moving globally left- or rightwards on a mid-grey background LCD projector (1024 x 768 Hz, 144 Hz) at a density of 2.6 dots/° and in a 5° diameter circular aperture centered at cartesian coordinates [−5°, 5°] (i.e., the bottom left quadrant of the visual field, relative to central fixation). Direction range of the dots was varied between 0° (total coherence) and 360° (complete random motion). The degree of difficulty was increased with improving task performance by increasing the range of dot directions within the stimulus. A 3:1 staircase design was implemented to allow us to compute a threshold level of performance for direction integration at the end of each timepoint (Das et al., 2014; Huxlin et al., 2009). For every 3 consecutive correct trials, direction range increased by 40°, while for every incorrect response, it decreased by 40°. The black dots making up the stimulus were 0.06° in diameter and moved at a speed of 10°/s over a time lapse of 250ms for a stimulus lifespan of 500ms. At every stimulus onset, an auditory beep was played for the subject. After each trial, auditory feedback indicated whether the response was correct or incorrect (see **Figure 2B** and **2C**).

### Transcranial Electrical Stimulation

Subjects were randomly assigned into 3 groups: In the first experimental group (n=17, 10 females), In-Phase (0° phase lag) bifocal tACS was applied over the right V1 and V5 areas. The second experimental group (n=18, 8 females), received Anti-Phase (180° phase lag) bifocal tACS over V1 and V5 areas, also in the right hemisphere. The control group (n=15, 6 females) received Sham (half cycle ramp-up) bifocal stimulation over identical V1 and V5 locations as the first two groups. The electrode placement on V1 and V5 were determined according to the 10-20 EEG system, i.e. over the O2 and P6 positions, respectively. Figure 1D gives an overview on the stimulating electrodes’ positions for the three groups.

Prior to the baseline recording, the Alpha peak frequency of each individual was determined over a 180s-long EEG resting-state recording with the eyes open, used thereafter as the individualized frequency for the tACS in time point TP0. Mean Alpha stimulation frequency for the In-Phase group was 9 Hz (range 7-11 Hz), for the Anti-Phase group:10 Hz (range 7-12 Hz) and for the Sham group: 10 Hz (range 7-11 Hz).

### Apparatus and devices

All experiments took place inside the same, shielded Faraday cage designed for EEG recordings, and under the same light conditions. Participants’ heads were placed over a chin-rest at a distance of 60 cm from the presentation screen, assuring a fixed position across all trials. The task ran on a Windows OS machine, based on a custom Matlab (The MathWorks Inc., USA) script, using the Psychophysics Toolbox.

Gaze and pupils’ movements were controlled in real time with an EyeLink 1000 Plus Eye Tracking System (SR Research Ltd., Canada) sampling at a frequency of 1000 Hz. The task required the subject to fixate a target at the center of the screen for every trial, with a maximal tolerance for eye deviation from this fixation target of about 1°. If the participant broke fixation during stimulus presentation, the moving stimulus froze and then disappeared, the trial was discontinued, and the computer played an unpleasant auditory tone. Once the participant repositioned their gaze correctly, a novel trial was started.

Bifocal tACS was delivered by means of two Neuroconn DC Plus stimulators (Neurocare group) triggered every cycle repeatedly to assure the chosen phase synchronization between the two stimulation sites. Custom-made, concentric, rubber electrodes of external diameter 5 cm, internal diameter of 1.5 cm and 2.5 cm of hole diameter were used to deliver stimulation. The intensity was fixed to 3mA corresponding to a current density of 0.18 mA/cm^2^. The electrodes were held by placing the EEG cap over them. The period of continuous stimulation, although it was slightly different for every participant, took on average ~13 ± 2 minutes (SEM), i.e. the time to complete 150 trials of the motion discrimination task described above.

EEG was recorded from a 64 channels passive system (Brain Products GMBH) at a sampling frequency of 5 kHz.

**Figure 1.**
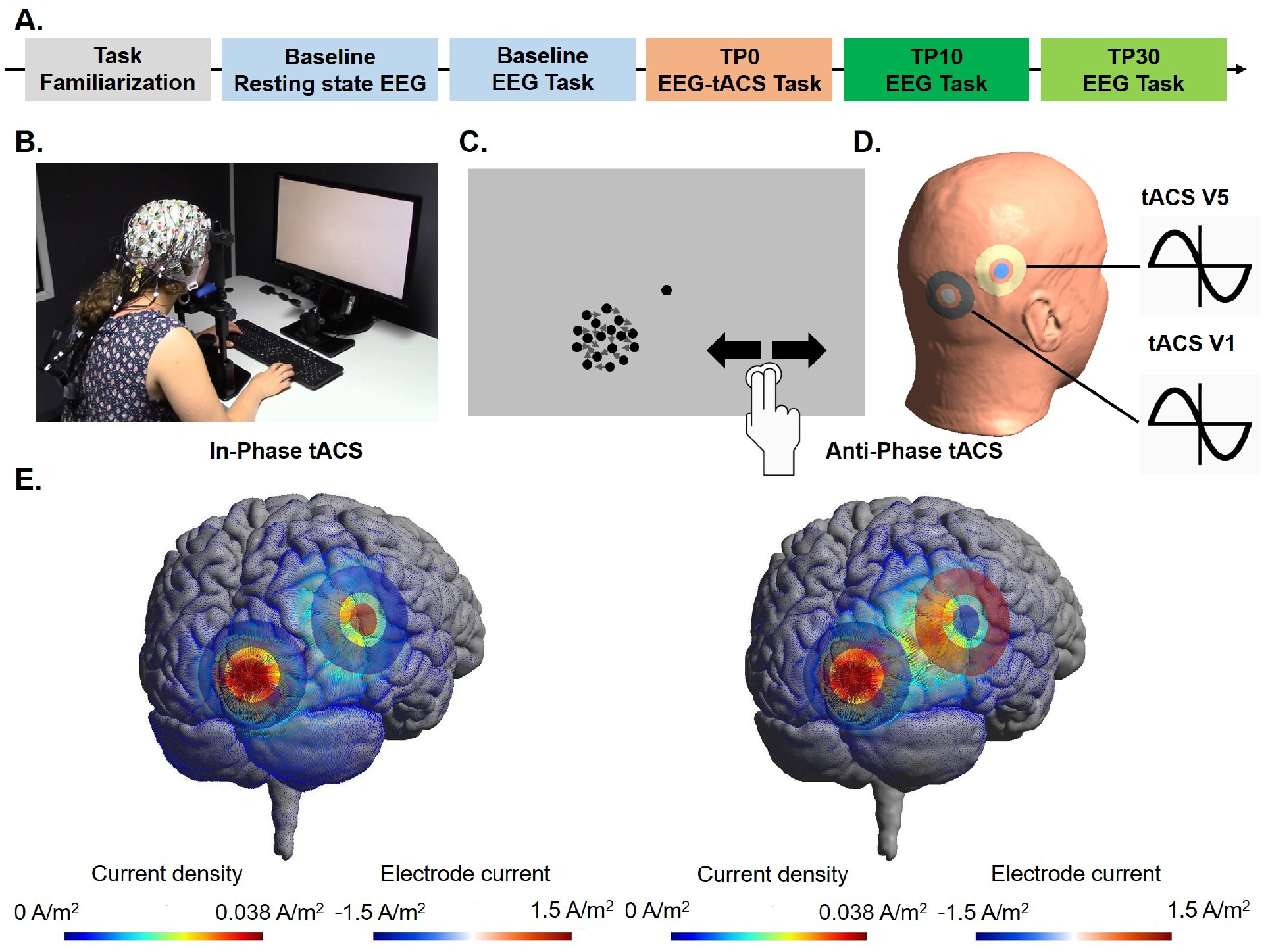
**(A) Experimental design**. The total duration of the experiment was around 3hrs. **(B) Real example**of the experimental setup inside the Faraday’s cage. The EEG system and an ongoing visual task are shown. **(B) Schematic example of the motion discrimination task**. **(C) Schematic of the bifocal tACS** applied with concentric electrodes over P6 and O2 while subject performs the global direction discrimination visual task. (**D) Electrical field 3D representation of bifocal tACS** at the two different phase differences (Thielscher et al., 2015). The dispersion of the field does not change over time in the two conditions, but rather the magnitude of the electrical field lines (Saturnino et al., 2017).

### Data Analysis

#### Behavioral data

For each subject and time point, we extracted direction range thresholds using all trials, by fitting a Weibull function, which defined the direction range level at which performance reached 75% correct. These direction range thresholds were then normalized to the maximum possible range of motion (360°), resulting in a normalized direction range threshold (NDR), a procedure previously described (Das et al., 2014; Huxlin et al., 2009).

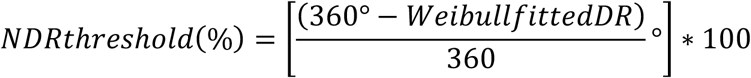

Finally, NDR thresholds were corrected for inter-individual variability in baseline performances by dividing all data by the individual baseline performances (referred as baseline-corrected NDR throughout the manuscript).

#### EEG data

All analyses were performed using MNE-Python (Gramfort et al., 2013) and customized scripts.

For the preprocessing, data were re-referenced to the average of signals, filtered through a Finite Response Filter of order 1, between 0.5 and 45Hz, epoched in 3s blocks. Every epoch corresponded to the time interval of a trial from the behavioral task. They were visually inspected to clear up noisy channels or unreadable trials. Bad channels were interpolated, data was re-sampled to 250Hz. Independent component analysis was used to remove physiological artifacts (i.e. eyeblinks, muscle torches).

For analyses in the frequency domain, Morlet wavelets convolution changing as a function of frequency was applied to 40 frequency bins, between 2 and 42Hz, increasing logarithmically.

For the source reconstruction analyses, data was re-referenced to the average of signals, noise covariance matrix was calculated to enhance the source approximation, a template brain and segmentation was used to compute the forward solution for 4098 sources per hemisphere. The inverse solution was calculated by means of MNE algorithm (Hämäläinen and Ilmoniemi, 1994). The points belonging to specific areas of interest (i.e. V1 and V5), were defined using the templates provided in the “SPM” open access database included in the MNE library (Wakeman and Henson, 2015). The source estimates were computed with dipole orientations perpendicular to the cortical surface (Lin et al., 2006). In order to extract one time-series per area of interest, we computed the first principal component from all source dipoles within each area. This first principal component is representing the source estimates associated with these pre-defined areas. Subsequently, a sign-flip was applied with the objective of avoiding sign ambiguities in the phase of different source estimates within the same area (Gramfort et al., 2012).

Specifically, the EEG metrics of interest computed were: Power Spectral Density (PSD) in the Alpha and Gamma band, both computed in the sensors’ space, Coherence in the Alpha and Gamma Band, V1 Alpha Phase to V5 Gamma Amplitude coupling (ZPAC-V1pV5a) and, V5 Alpha Phase to V1 Gamma Amplitude coupling (ZPAC-V5pV1a), computed in the sources’ space. All these variables were baseline-normalized. Moreover, the Phase Amplitude coupling (i.e. PAC) was standardized to avoid confounders by creating a non-parametrized distribution of values to which to compare the observations through a Z-score transformation (i.e. ZPAC) (Canolty et al., 2006; Cohen, 2014).

Thus, PSD (Φ) was calculated taking an average of all electrodes through the Welch’s estimator (Welch, 1967), that considers averaging PSDs from different windows, according to the formula:

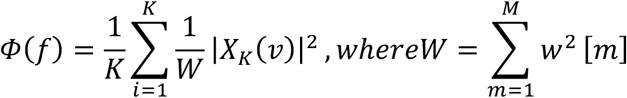

Where K corresponds to the number of segments where a windowed Discret Fourier Transform is computed, X is the segment where it is computed at some frequency v and w is the window segment

(Magnitude-square) Coherence (Carter, 1987) was calculated through:

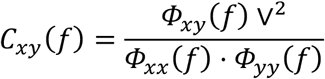

V1-V5 coherence analyses are used to investigate frequency-specific phase coupling between these source areas. Although coherence values might be biased due to source leakage effects (Palva et al., 2018), we included this metric because it is of relevance given our brain stimulation approach.

Phase Amplitude coupling (PAC) (Canolty et al., 2006) was obtained through:

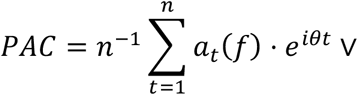

Where *t* corresponds to a certain time point, *a* denotes the power at a certain specific frequency for this specific time point, *i* is the imaginary variable, θ the phase angle and *n* the number of time points. In the manuscript, we will refer to ZPAC V1 Alpha phase – V5 Gamma amplitude (ZPAC-V1p_Alpha_V5a_Gamma_) as a bottom-up modulation and PAC V1 Gamma amplitude – V5 Alpha phase (ZPAC-V1a_Gamma_V5p_Alpha_) as a top-down modulation (see (Nandi et al., 2019)). In order to verify the lack of influence concerning the signal leakage problem in the calculation of the Phase Amplitude Coupling, computations showing the modulation of the phase and amplitude within the same areas of source estimates were computed (See supplementary figure 2).

### Statistical Analyses

#### Behavior

Statistical analyses were carried out using mixed-effect linear models. The evolution of the baseline-corrected NDR was investigated as a dependent variable, with stimulation group and time points as the main fixed effects.

#### EEG metrics

PSD (Gamma and Alpha components across time) significance within subjects was tested through a sliding FDR-corrected T-test. Significance within subjects in the Coherence and Phase-Amplitude coupling spectrums were evaluated through non-parametric permutation tests and clusters-based corrected for multiple comparisons. Differences were considered significant when p < 0.05.

A mixed linear model was performed in order to evaluate the variability of the chosen EEG metric (dependent variable) over time, among stimulation groups.

#### Best EEG metric

In order to determine the EEG metric that had the highest impact on the behavioral scores and then reduce the model space of the baseline-corrected NDR mixed linear model, an embedded regularization method (i.e., least absolute shrinkage and selection operator - Lasso) was applied (Tibshirani, 1996) following the Langragian version of the formula:

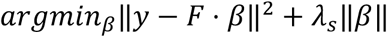

Where β corresponds to the unknown vector of weighted coefficients estimated for every metric (regression coefficient), y is the matrix with all the labeled metrics, λ is in charge of the variable selection and F correspond to the acquired data points. Lasso was selected due to the fact that it provides a preferred solution with the highest sparsity given the shrink provided by the penalty term. The vector of λ chosen consisted in 30 testing points spaced between 0 and 1. The number of iterations was set to 1000.

#### Behavior + EEG

As a second step, covariates that could explain variance in NDR outcome and a possible interaction effect with stimulation group were added to the first mixed linear model. A random intercept per subject was used to correct for the dependency between time points for all models. The residuals of each statistical model were tested for normality by inspecting histograms and through the omnibus normality test (D’Agostino and Pearson, 1973).

## Supporting information

SOM

## ACKNOWLEDGEMENTS

This research was funded by the Bertarelli Foundation (Catalyst BC77O7 to FCH & ER), by the Swiss National Science Foundation (PRIMA PR00P3_179867 to ER) and by the Defitech Foundation (to FCH).

## AUTHOR CONTRIBUTIONS

F.C.H. and E.R. developed the research idea and F.C.H., R.S.G., E.R., K.H. the experimental design. R.S.G. and E.R. were in charge of the data acquisition and data analyses. S.Z., M.S. added to the analyses. R.S.G. drafted first version of the manuscript. All authors revised the manuscript significantly. F.C.H. and E.R provided the funding.

## COMPETING INTERESTS

The authors declare no competing interests.

## Notes

### Competing Interest Statement

The authors have declared no competing interest.

