## Supplementary material for "Bifocal tACS Enhances Visual Motion Discrimination by Modulating Phase Amplitude Coupling Between V1 and V5 Regions": SOM

**SUPPLEMENTARY MATERIALS**

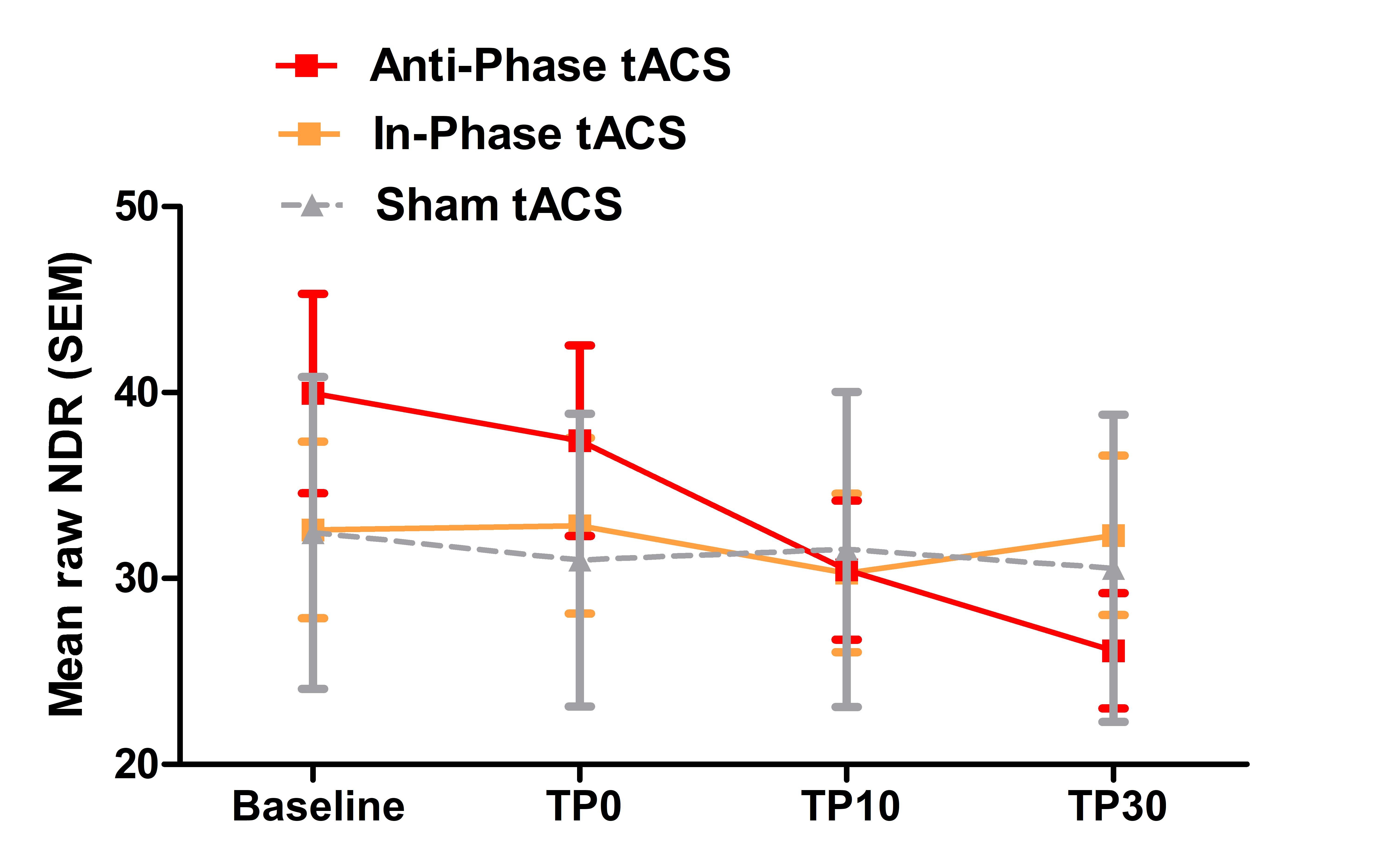

**Supplementary Figure 1. NDR (Normalized Direction Range) threshold evolution across time-points for the three stimulation conditions.** Bars correspond to Standard Errors of the Mean (SEM).

|  | **Baseline** | **TP0** | **TP10** | **TP30** |
| --- | --- | --- | --- | --- |
| **In-Phase** | 32.61 (4.75) | 32.84 (4.73) | 30.30 (4.27) | 32.32 (4.29) |
| **Anti-Phase** | 39.95 (5.36) | 37.42 (5.13) | 30.45 (3.73) | 26.11 (3.11) |
| **Sham** | 32.45 (8.39) | 30.99 (7.87) | 31.56 (8.48) | 30.55 (8.27) |

**Supplementary Table 1. Mean raw NDR (Normalized Direction Ratio) values (SEM) for the three groups and four timepoints (TP).** The group differences, although not significant (see the Results section) at baseline and the large overall inter-individual variability prompted us to perform a baseline-correction procedure by normalizing (dividing) all performances by the individual baseline value (see the Methods section).

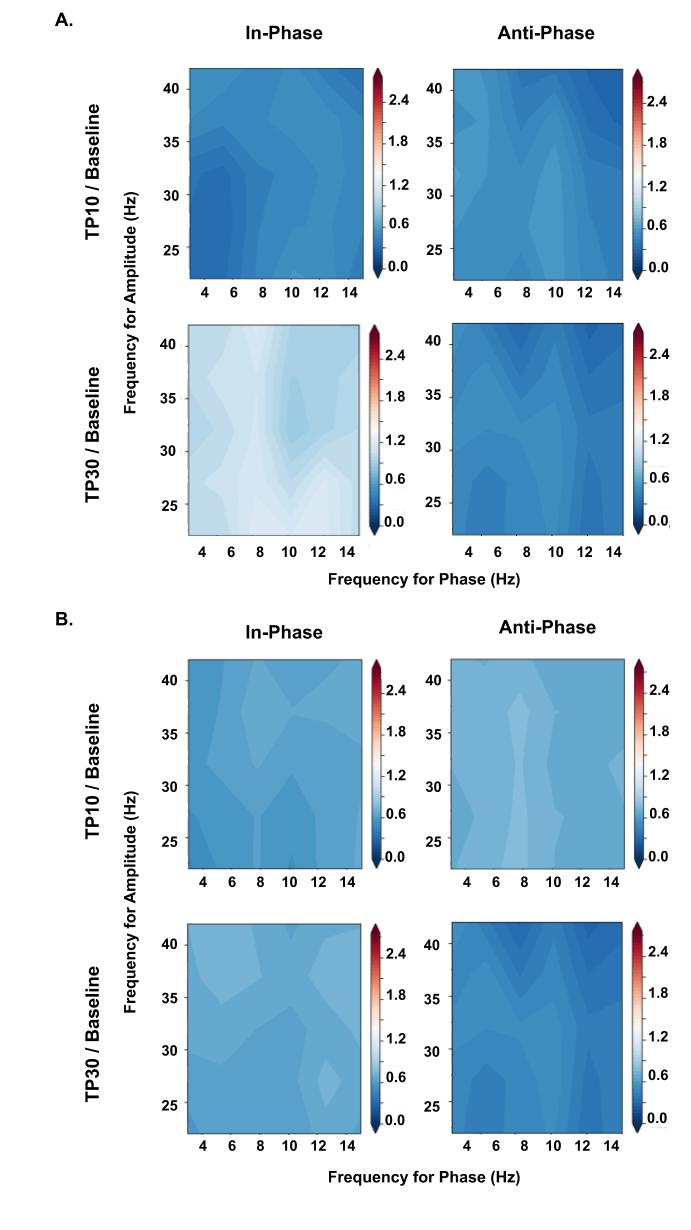

**Supplementary Figure 2.** Alpha Phase - Gamma Amplitude cross frequency spectrums evaluated within the same brain area for the two conditions shown to be significantly different from each other (i.e. In-Phase vs. Anti-Phase). There is no Alpha-Gamma modulation of interest within the areas.  **A.** **V1 Alpha Phase – V1 Gamma Amplitude B. V5 Alpha Phase – V5 Gamma Amplitude**

|  | **Coef.** | **Std.Err.** | **z** | **P>\|z\|** | **[0.025** | **0.975]** |
| --- | --- | --- | --- | --- | --- | --- |
| **TP10-TP0** | -0.05 | 0.038 | -1.314 | 0.189 | -0.124 | 0.024 |
| **TP30-TP0** | -0.067 | 0.038 | -1.758 | 0.079 | -0.141 | 0.008 |
| **TP30-TP10** | -0.017 | 0.038 | -0.443 | 0.657 | -0.091 | 0.057 |
| **Anti-Phase-In-Phase (TP0)** | -0.257 | 0.106 | -2.43 | 0.015 | -0.464 | -0.05 |
| **Sham-In-Phase (TP0)** | -0.16 | 0.102 | -1.564 | 0.118 | -0.36 | 0.04 |
| **Sham-Anti-Phase (TP0)** | 0.097 | 0.104 | 0.936 | 0.349 | -0.107 | 0.301 |
| **Anti-Phase-In-Phase (TP10)** | -0.257 | 0.106 | -2.43 | 0.015 | -0.464 | -0.05 |
| **Sham-In-Phase (TP10)** | -0.16 | 0.102 | -1.564 | 0.118 | -0.36 | 0.04 |
| **Sham-Anti-Phase (TP10)** | 0.097 | 0.104 | 0.936 | 0.349 | -0.107 | 0.301 |
| **Anti-Phase-In-Phase (TP30)** | -0.257 | 0.106 | -2.43 | 0.015 | -0.464 | -0.05 |
| **Sham-In-Phase (TP30)** | -0.16 | 0.102 | -1.564 | 0.118 | -0.36 | 0.04 |
| **Sham-Anti-Phase (TP30)** | 0.097 | 0.104 | 0.936 | 0.349 | -0.107 | 0.301 |
| **ZPAC_V1pV5a** | 0.015 | 0.012 | 1.292 | 0.196 | -0.008 | 0.039 |
| **ZPAC_V1pV5a : Anti-Phase (In-Phase + TP10)** | -0.071 | 0.036 | -1.975 | 0.048 | -0.142 | -0.001 |
| **ZPAC_V1pV5a : Sham (In-Phase + TP10)** | -0.023 | 0.03 | -0.771 | 0.44 | -0.081 | 0.035 |
| **ZPAC_V1pV5a : Sham (Anti-Phase + TP10)** | 0.048 | 0.029 | 1.667 | 0.095 | -0.008 | 0.105 |
| **ZPAC_V1pV5a : Anti-Phase (In-Phase + TP30)** | -0.071 | 0.036 | -1.975 | 0.048 | -0.142 | -0.001 |
| **ZPAC_V1pV5a : Sham (In-Phase + TP30)** | -0.023 | 0.03 | -0.771 | 0.44 | -0.081 | 0.035 |
| **ZPAC_V1pV5a : Sham (Anti-Phase + TP30)** | 0.048 | 0.029 | 1.667 | 0.095 | -0.008 | 0.105 |
| **ZPAC_V5pV1a** | -0.007 | 0.011 | -0.629 | 0.53 | -0.029 | 0.015 |
| **ZPAC_V5pV1a : Anti-Phase (In-Phase + TP10)** | -0.055 | 0.069 | -0.786 | 0.432 | -0.191 | 0.082 |
| **ZPAC_V5pV1a : Sham (In-Phase + TP10)** | 0.006 | 0.049 | 0.115 | 0.908 | -0.09 | 0.101 |
| **ZPAC_V5pV1a : Sham (Anti-Phase + TP10)** | 0.06 | 0.051 | 1.189 | 0.234 | -0.039 | 0.16 |
| **ZPAC_V5pV1a : Anti-Phase (In-Phase + TP30)** | -0.055 | 0.069 | -0.786 | 0.432 | -0.191 | 0.082 |
| **ZPAC_V5pV1a : Sham (In-Phase + TP30)** | 0.006 | 0.049 | 0.115 | 0.908 | -0.09 | 0.101 |
| **ZPAC_V5pV1a : Sham (Anti-Phase + TP30)** | 0.06 | 0.051 | 1.189 | 0.234 | -0.039 | 0.16 |

**Supplementary Table 2 - NDR Mixed Linear Model with all the independent variable contributions.** Beta coefficients, P-values and Confidence intervals are presented for all the comparisons and interactions among time points, stimulation groups and the EEG markers: V1 Alpha Phase V5 Gamma Amplitude coupling (ZPAC_V1pV5a) and V1 Gamma Amplitude V5 Alpha Phase coupling (ZPAC_V5pV1a).

|  | **Coef.** | **Std.Err.** | **z** | **P>\|z\|** | **[0.025** | **0.975]** |
| --- | --- | --- | --- | --- | --- | --- |
| **TP30-TP10** | -0.769 | 0.402 | -1.915 | 0.055 | -1.556 | 0.018 |
| **Anti-Phase-In-Phase (TP10)** | -0.836 | 0.894 | -0.935 | 0.35 | -2.588 | 0.916 |
| **Sham-In-Phase (TP10)** | 0.173 | 0.859 | 0.202 | 0.84 | -1.51 | 1.856 |
| **Sham-Anti-Phase (TP10)** | 1.009 | 0.876 | 1.152 | 0.249 | -0.708 | 2.726 |
| **Anti-Phase-In-Phase (TP30)** | -0.836 | 0.894 | -0.935 | 0.35 | -2.588 | 0.916 |
| **Sham-In-Phase (TP30)** | 0.173 | 0.859 | 0.202 | 0.84 | -1.51 | 1.856 |
| **Sham-Anti-Phase (TP30)** | 1.009 | 0.876 | 1.152 | 0.249 | -0.708 | 2.726 |

**Supplementary Table 3 - ZPAC_V1pV5a Mixed Linear Model.** Beta coefficients, P-values and Confidence intervals are presented for all the comparisons and interactions among time points and stimulation groups

|  | **Coef.** | **Std.Err.** | **z** | **P>\|z\|** | **[0.025** | **0.975]** |
| --- | --- | --- | --- | --- | --- | --- |
| **TP30-TP10** | 0.409 | 0.383 | 1.066 | 0.286 | -0.343 | 1.161 |
| **Anti-Phase-In-Phase (TP10)** | -0.695 | 1.045 | -0.665 | 0.506 | -2.744 | 1.353 |
| **Sham-In-Phase (TP10)** | -1.413 | 1.009 | -1.401 | 0.161 | -3.39 | 0.564 |
| **Sham-Anti-Phase (TP10)** | -0.718 | 1.025 | -0.7 | 0.484 | -2.727 | 1.292 |
| **Anti-Phase-In-Phase (TP30)** | -0.695 | 1.045 | -0.665 | 0.506 | -2.744 | 1.353 |
| **Sham-In-Phase (TP30)** | -1.413 | 1.009 | -1.401 | 0.161 | -3.39 | 0.564 |
| **Sham-Anti-Phase (TP30)** | -0.718 | 1.025 | -0.7 | 0.484 | -2.727 | 1.292 |

**Supplementary Table 4 - ZPAC_V5pV1a Mixed Linear Model.** Beta coefficients, P-values and Confidence intervals are presented for all the comparisons and interactions among time points and stimulation groups
